# Carrion fly-derived DNA metabarcoding is an effective tool for mammal surveys: evidence from a known tropical mammal community

**DOI:** 10.1101/133405

**Authors:** Torrey W. Rodgers, Charles C. Y. Xu, Jacalyn Giacalone, Karen M. Kapheim, Kristin Saltonstall, Marta Vargas, Douglas W. Yu, Panu Somervuo, W. Owen McMillan, Patrick A. Jansen

## Abstract

Metabarcoding of vertebrate DNA derived from carrion flies has been proposed as a promising tool for biodiversity monitoring. To evaluate its efficacy, we conducted metabarcoding surveys of carrion flies on Barro Colorado Island (BCI), Panama, which has a well-known mammal community, and compared our results against diurnal transect counts and camera-trapping. We collected 1084 flies in 29 sampling days, which were pooled into 102 DNA extractions. We then conducted metabarcoding with mammal-specific (16S) and vertebrate-specific (12S) primers targeting mtDNA, and sequenced these amplicons on Illumina MiSeq. For taxonomic assignment, we compared BLAST with the new program PROTAX, and we found PROTAX significantly improved species identifications. We detected 20 mammal, four bird, and one lizard species from carrion fly metabarcoding, all but one of which were previously known from BCI. Fly metabarcoding detected more mammal species than concurrent transect counts (29 sampling days, 13 species) and concurrent camera-trapping (84 sampling days, 17 species), and detected 67% of the number of mammal species documented by 8 years of transect counts and camera-trapping combined, although fly metabarcoding missed several abundant species. This study demonstrates that carrion fly metabarcoding is a powerful tool for mammal biodiversity surveys, and has the potential to detect a broader range of species than more commonly used methods.

## Introduction

Due to the rapid decline in biodiversity worldwide, there is urgent need for more efficient techniques to survey and monitor biodiversity. Traditional surveys based on direct observations are costly and time consuming, results can vary an unknown degree across observers, and rare or cryptic species are often overlooked. Camera trapping has emerged as a more efficient survey method, especially for large-bodied vertebrates (Beaudrot et al. 2016), but it has limited ability to detect small-bodied, arboreal, and volant species. Moreover, camera traps are expensive for use at large scales and are subject to damage and theft. An efficient technique capable of detecting a wide range of vertebrate species with limited bias would greatly increase research’s ability to monitor vertebrate diversity, and to gauge the effectiveness of conservation measures such as nature reserves.

One such potential method has emerged from the field of environmental DNA (eDNA) sampling, which uses trace amounts of DNA from the environment for species detection (Bohmann et al. 2014). This approach, particularly when used in conjunction with next-generation metabarcode sequencing, shows great potential for efficient monitoring of biodiversity (Ji et al. 2013; Taberlet et al. 2012; Yu et al. 2012). To date, the majority of vertebrate eDNA research has focused on aquatic species because eDNA is easy to collect from water (Thomsen et al. 2012). Collecting eDNA from terrestrial vertebrates is more difficult. Researchers have used soil (Andersen et al. 2012), browsed twigs (Nichols et al. 2012), prey carcasses (Wheat et al. 2016), and drinking water (Rodgers & Mock 2015) for eDNA detection of terrestrial species, but these approaches have limited scope. So far, the most promising approach for use across a broad range of species is mass trapping and metabarcoding of invertebrates that feed on vertebrates (Bohmann et al. 2013; Calvignac-Spencer et al. 2013a). Such invertebrate ‘samplers’ tested to date include leeches (Schnell et al. 2015), mosquitos (Logue et al. 2016), dung beetles (Gillett et al. 2016) and carrion flies (Calvignac-Spencer et al. 2013b; Lee et al. 2015; Schubert et al. 2014).

Carrion flies are species from the families Calliphoridae and Sarcophagidae that feed and oviposit upon dead animals, open wounds, and feces. When carrion flies feed, they ingest vertebrate DNA. Carrion flies are ideal candidates for eDNA surveys because they are easy to trap, are ubiquitous worldwide, and are believed to feed opportunistically on vertebrates of all sizes, including terrestrial species, volant species, and species occupying the forest canopy. In a remarkable study, Calvignac-Spencer et al. (2013b) used Sanger sequencing to detect 20 different mammal species from just 115 flies collected in Côte d’Ivoire and Madagascar. They also used Roche GS FLX (‘454’) pyrosequencing to metabarcode a subset of samples, and were able to detect the majority of species that had been detected with Sanger sequencing, plus several others.

Although carrion fly metabarcoding is clearly a promising method, there is still a need to quantify its effectiveness relative to conventional methods. There is also a need to test new methods for assigning taxonomies to sequence data. This step may be particularly error prone in eDNA studies, because target amplicons for eDNA are short due to the need to amplify degraded DNA, and thus have lower taxonomic information content. Because reference sequence databases are often incomplete, taxonomic assignment software has a tendency toward overconfidence, meaning that an OTU sequence may be assigned to a species that is in the database and fairly similar, when the true species is not in the database (Somervuo et al. 2017). Correct species-level identification is especially important when applied to vertebrates, as incorrect assignment of an OTU to an endangered species, as opposed to a less endangered congener (or vice versa) can have a large impact on conservation decision making.

With these goals in mind, we conducted a field test on Barro Colorado Island (BCI) in Panama, where the vertebrate fauna is well documented, particularly for mammals. We collected carrion flies for metabarcoding, and we compared results with datasets from annual, diurnal transect counts (distance sampling) and semi-continuous camera-trapping of the mammal community for 8 years leading up to, and concurrent with, fly collection. In addition, we compared taxonomic assignments between the most commonly used method, BLAST (Altschul et al. 1990) and the new method PROTAX (Somervuo et al. 2017), which uniquely takes into account the possibility that an OTU sequence belongs to a species that is not in the reference database, thus avoiding overconfident assignments. We find that metabarcoding of carrion flies is an effective, but imperfect, method for surveying mammal communities, and use of PROTAX substantially improves taxonomic assignment.

## Materials and Methods

### Study site

Field work was conducted on Barro Colorado Island, a 1,560-ha island in the Panama Canal waterway (Fig. 1). BCI (9°10’N, 79°51’W) sits within Gatun Lake, an artificial body of water created in 1912 by the damming of the Chagres River to create the Panama Canal, and is part of the protected 54-km^2^ Barro Colorado Nature Monument. BCI has 108 known mammal species including 74 bats, and 34 non-volant species, (Glanz 1982 and current expert information). However some of these species such as jaguar (*Panthera onca*), puma (*Puma concolor*), and jaguarundi (Herpailurus yagouaroundi) are infrequent, non-resident visitors to the island.

**Figure 1.**
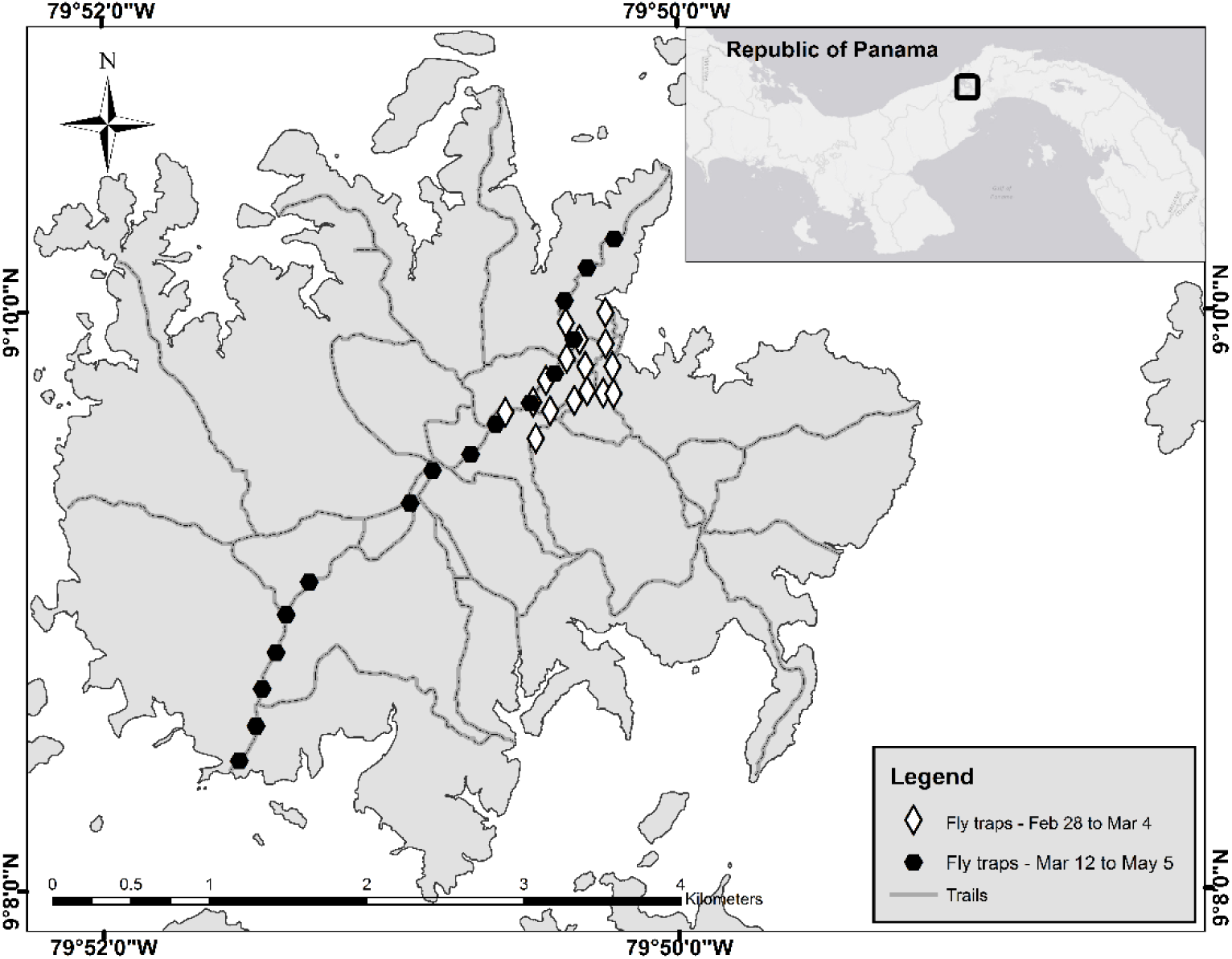
Location of carrion-fly trapping on Barro Colorado Island, Panama.

### Fly collection

Flies were collected between 10 Feb and 5 May 2015, in three trapping sessions. First, flies were collected opportunistically from 10–23 Feb using a variety of trap types within 1000 m of the labs on BCI, to determine the best methods for sampling. Based on results from this session, we used a trap type modified from www.blowflies.net/collecting.htm, for all subsequent trapping. Traps were baited with raw pork hung in a cup surrounded by fine cloth netting to keep flies from landing directly on the bait, and bait was replaced every 2-3 days. Second, from 28 Feb–4 Mar, traps were placed along trails at 16 trap locations, each containing 2 traps, spaced roughly 200 m apart in a non-uniform grid (Fig 1). Third, from 12 Mar to 5 May 2015, flies were collected from traps placed in a transect crossing the island with 16 trap locations, each containing 2 traps, placed every 250 m along a trial in a roughly straight line (Fig 1), for 10 sampling days. In all three collection efforts, flies were removed from traps once or twice daily and placed in a −40 °C freezer within 2 hours of collection, because DNA degradation causes detection success to decline 24 hours after fly feeding (Lee et al. 2015).

### Library preparation and sequencing

DNA was extracted from flies using the GeneMATRIX Stool DNA Purification Kit (Roboklon, Berlin, Germany). To reduce extraction costs, up to 16 flies were pooled for each extraction. Flies were first cut into several pieces with sterile scissors and placed into 2.5ml Polypropylene 96 Deep Well Plates along with stainless steel, 5/32” grinding balls (OPS diagnostics; Lebanon, NJ). Up to 4 flies and 100 μl of lysis buffer per fly were added to each well. Plates were shaken on a TissueLyser II (Qiagen; Germantown, MD) until fly tissue was homogenized in the lysis buffer. An equal volume of the resulting homogenate per fly was then pooled in a total volume of 160 μl, and added to the extraction kit bead tube.

Extractions then proceeded following manufacturer’s recommendations. For flies collected in the first trapping session, each extraction included 16 flies. For flies collected in the second and third trapping session, each extraction contained all flies collected at the same trap location on the same day, up to a maximum of 16 flies. To examine if pooling had an effect on species detection, 24 flies were also homogenized and extracted individually, and 20 μl of homogenate from each of these same 24 flies was also pooled and extracted in 2 samples containing 12 flies each. All extractions included blank controls that were included in the PCR step to test for contamination.

PCR was performed on all samples using a mammal-specific primer set (16Smam1, 16Smam2; Boessenkool et al. 2012) targeting 130-138 bp of the mitochondrial 16S rRNA locus, and a pan-vertebrate primer set (12SV5F, 12SV5F; Riaz et al. 2011) targeting 140-143 bp of the mitochondrial 12S locus. Reactions also included human blocking primers (16Smam_blkhum3; Boessenkool et al. 2012; 12S_V5_blkhum; Calvignac-Spencer et al. 2013b) and *Sus* blocking primers (16Smam_blkpig, 12S_V5_blkpig; Calvignac-Spencer et al. 2013b) to decrease competition from contaminating DNA from pork bait. Primers also included a 5’ addition of a 33bp Illumina-specific sequence for addition of illumina adapters in a second round of PCR. A minimum of two PCR replicates per sample were performed in 10 μl volumes. For 16S, reactions included 0.2 um of each primer, 1 μm of human blocking primer (5x) 4 μm of *Sus* blocking primer (20X), 200 μm dNTP, 4 mM MgCl2, 1X PCR buffer, 1.25 U Platinum® Taq polymerase (Invitrogen) and 3 μl of template DNA. For 12S, volumes were the same, except MgCl_2_ was reduced to 2.5 mM. Cycling conditions were 10 min at 95 °C, followed by 42 cycles (16S) or 47 cycles (12S) of 30 s at 95 °C, 30 s at 64 °C, and 1 min at 72 °C, with a final extension of 10 min at 72 °C. Following the initial PCR, sample replicates were pooled, and a second PCR was conducted to add Illumina flow cell binding sequences, and unique 8bp sample specific indexes to each end of each amplicon. Reactions included 1 μl each of the forward and reverse illumina index tags, 2.5 ul of 10X Qtaq buffer, 1.5 Mm MgCl2, and 1.25 units of Qiagen Qtaq. Cycling conditions were 3 min at 94°C, followed by 6 cycles of 45 s at 94 °C, 60 s at 50 °C, and 1 min at 72 °C, with a final extension of 10 min at 72 °C. All PCRs included no-template negative controls to test for contamination. Samples were purified and normalized using SequalPrep™ Normalization Plates (Thermo-Fisher Scientific; Waltham, MA). All samples were pooled in equimolar concentration, and pools were concentrated using standard Ampure XP beads (Beckman Coulter; Indianapolis, IN). The concentrated pool was quantified using a Qubit fluorometer and quality checked using a Bioanalyzer High Sensitivity Kit. The library was sequenced on an Illumina MiSeq using a 2 x 250 PE sequencing kit.

### Sequence filtering

After demultiplexing, paired ends were merged for each sample using PEAR v. 0.9.6 (Zhang et al. 2014) with default parameters, except minimum overlap (-v) = 100, minimum length (-n) = 100, and quality score threshold (-q) = 15. We then separated 12S and 16S sequences and removed primers and remaining adapters with *cutadapt* v. 1.3 (Martin 2011), keeping sequences with a minimum length (-m) of 10. (Sequences had been previously trimmed to minimum 100. Mean ± 1 s.e. length after trimming was 109.78 ± 0.13 for 12S and 93.70 ± 0.08 for 16s.) Sequences were subsequently pooled and filtered for chimeras with the identify_chimeric_seqs.py script and the usearch61 method within the QIIME environment (Caporaso et al. 2010; Edgar 2010). We used the pick_de_novo_otus.py script with the uclust method to cluster sequences into OTUs (Operational Taxonomic Units) at 97% similarity, create a table of OTU frequency in each sample, and pick representative sequences (Caporaso et al. 2010; Edgar 2010). We removed OTUs with fewer than 20 sequences using the filter_otus_from_otu_table.py script (Caporaso et al. 2010). To remove OTUs from contaminants such as pork bait, human DNA, and bacteria, we conducted an initial BLAST search and removed all OTUs with a top hit for *Sus*, *Homo*, or bacteria. For 12S we used this initial BLAST search to separate OTUs into mammal, bird and reptile groups for downstream taxonomic assignment.

### Taxonomic assignment

For mammalian taxonomic assignment, we compared two methods: the most commonly used method BLAST (Altschul et al. 1990) and the new probabilistic taxonomic placement method PROTAX (Somervuo et al. 2017). For non-mammals, only BLAST was used. BLAST searches were conducted against the National Center for Biotechnology Information (NCBI) non-redundant database with default settings except for e-value ≤ 1e-6, and percent identity ≥ 95%. We processed the BLAST output in two ways for each OTU: selecting the top hit, and also using the lowest common ancestor (LCA) algorithm in MEGAN 5.11.3 (Huson et al. 2007; default settings except top percent = 5).

For PROTAX, we generated probabilities of taxonomic placement for each OTU at 4 taxonomic ranks (order, family, genus, and species) (Somervuo et al. 2016). Unlike other taxonomic assignment software, PROTAX is a statistical wrapper that processes the output of one or more other taxonomic assignment methods and takes into account the uncertainty of taxonomic assignment caused by species that do not have reference sequences. Briefly, this is achieved by training a model against a reference dataset that includes all available reference sequences for the taxon of interest, plus a full Linnaean taxonomy for that taxon which includes all named species, including those that do not have reference sequences. By taking into account the taxonomic information content of both the gene sequence being assigned and the reference sequence database, PROTAX removes the inherent ‘over-assignment’ bias of other taxonomic assignment software. This makes PROTAX especially suited for markers that have highly incomplete reference databases, such as 16S and 12S.

For the reference sequence databases, we downloaded all available mammalian mitochondrial 16S and 12S sequences from GenBank, truncated each species to a maximum of 10 sequences, removed ambiguous bases, and used ecoPCR within OBITools 1.2.6 (Ficetola et al. 2010) to extract the amplicon regions used in this study (Clark et al. 2016). This resulted in 5243 16S sequences representing 1733 species and 5395 12S sequences representing 1935 species. For the taxonomic database, we downloaded full ranks for the 6711 species in the NCBI Mammalia taxonomy. We are aware that the NCBI taxonomy is not complete for the Mammalia, but the important benefit is that the names are consistent between the sequence and taxonomic databases, and even this incomplete Mammalia taxonomy has ca. 3.8X and 3.4X the number of species as represented in the 16S and 12S reference-sequence databases, respectively, which allows us to contrast PROTAX with BLAST.

To start, all OTU representative sequences were pairwise compared to the reference sequences using LAST version 744 (Kielbasa et al. 2011), and the PROTAX model used the maximum and second-best similarities. We parameterized three PROTAX models, one unweighted model in which all mammal species were given an equal prior weight of being in the sampling location, and two weighted models in which species independently known to be present in Panama (Panama-weighted) or BCI (BCI-weighted) were given a 90% prior probability of being in the sampling location, with all other species being given a prior probability of 10%. The accuracy and bias of the trained PROTAX model at each taxonomic rank were estimated by plotting the cumulative predicted probabilities against the cumulative number of cases in which the outcome with the highest probability was correct when training the model.

For all methods (BLAST top hit, BLAST plus MEGAN, and PROTAX unweighted and weighted), we estimated accuracy by calculating the percentage of OTUs assigned to a genus or species known from BCI (rate of correct assignment) and the percentage of OTUs assigned to a genus or species known to not be present on BCI (rate of clear false positives). We also counted the number of mammal species correctly identified with each method. Given that we only used wild-caught flies, we cannot directly test the probability that an OTU was assigned incorrectly to a species known from BCI. However, the PROTAX output does give us a probability of correct assignment for each OTU at each rank.

### Transect counts and camera trapping

We compared results from fly metabarcoding with results of two traditional survey methods. First, diurnal mammal transect counts were carried out yearly from 2008-2015 during January and/or February of each year. Transects covered all trails on BCI (Fig 1) each year, with a mean distance of 120km walked per year. An average walking rate of 1km/hr was maintained. Record of each mammal sighting included date, time, location, species, and number of individuals. All censuses were conducted between 06:40 and 12:00, and were thus unlikely to detect nocturnal animals. In 2015, the year fly sampling was conducted, 151.1 km of transect was surveyed from 24 Jan to 22 Feb.

Second, camera traps were operated continuously across BCI from 2008-2015. A mean of 25 (range 19-34) trail cameras (PC900 and RC55 Reconyx Inc., Holmen, Wisconsin) were mounted at knee height, with a spacing of 500-1000 meters. Thus, they were unlikely to capture arboreal or volant species. Additional cameras were deployed for shorter periods. Cameras were checked at least once every 6–7 months, and replaced or repaired if no longer functioning. This effort resulted in a total of 39,151 total camera-days. For the period concurrent with fly collection, Feb 10 to May 5 2015, 26 cameras were active for the entire period, resulting in a total effort of 1,967 camera-days.

### Methods comparison

We compared the total number of mammal species detected by fly metabarcoding with the total number of species detected by concurrent (2015) and long term (2008-2015) transect counts and camera trapping. In addition, we fit species accumulation curves for each of the 2015 surveys using the *specaccum* function in the R package *vegan* 2.4-1 (Oksanen et al. 2016) with *method* = random and 10,000 permutations.

## Results

### Transect counts and camera trapping

In the 2015 transect counts, we detected 13 mammal species, whereas in 2008-2015 transect counts, we detected 17 mammal species. With concurrent 2015 camera trapping, we detected 17 mammal species, whereas in 2008-2015 camera trapping, we detected 26 mammal species. Species detected by the two methods overlapped only partially, so that the 2015 datasets combined detected a total of 22 mammal species, and the 2008-2015 datasets combined detected a total of 30 mammal species.

### Fly metabarcoding

#### Fly collection, extraction and sequencing

A total of 1084 flies were collected and pooled into 102 pooled samples and 24 single-fly samples. We obtained 2,780,574 initial reads, which were reduced to 2,288,009 after quality filtering (1,504,440 16S; 783,569 12S). OTU clustering, removal of OTUs with fewer than 20 reads, and removal of OTUs assigned to human, *Sus* (pork bait), and bacteria resulted in a final set of 54 OTUs for 16S and 63 OTUs for 12S (49 mammal, 12 bird, and 2 reptile OTUs).

#### Mammal species detection

The number of mammal species detected differed between assignment methods (Fig 2a). BCI-weighted PROTAX resulted in the detection of 20 total mammal species (table 1), the most of any method (16 species from 53 OTUs with 16S, 13 species from 37 OTUs with 12S, and 9 species with both markers). Assignment of multiple OTUs to the same species is expected in metabarcoding since the sequence-clustering step typically applies a single similarity threshold across all OTUs, which can result in species splitting. Fifteen of these 20 species had PROTAX probabilities of > 0.9, but several had somewhat low PROTAX probabilities despite their known presence on BCI (table 1). Panama-weighted and unweighted PROTAX resulted in detection of 19 total mammal species (13 with 16S and 12 with 12S).

**Figure 2.**
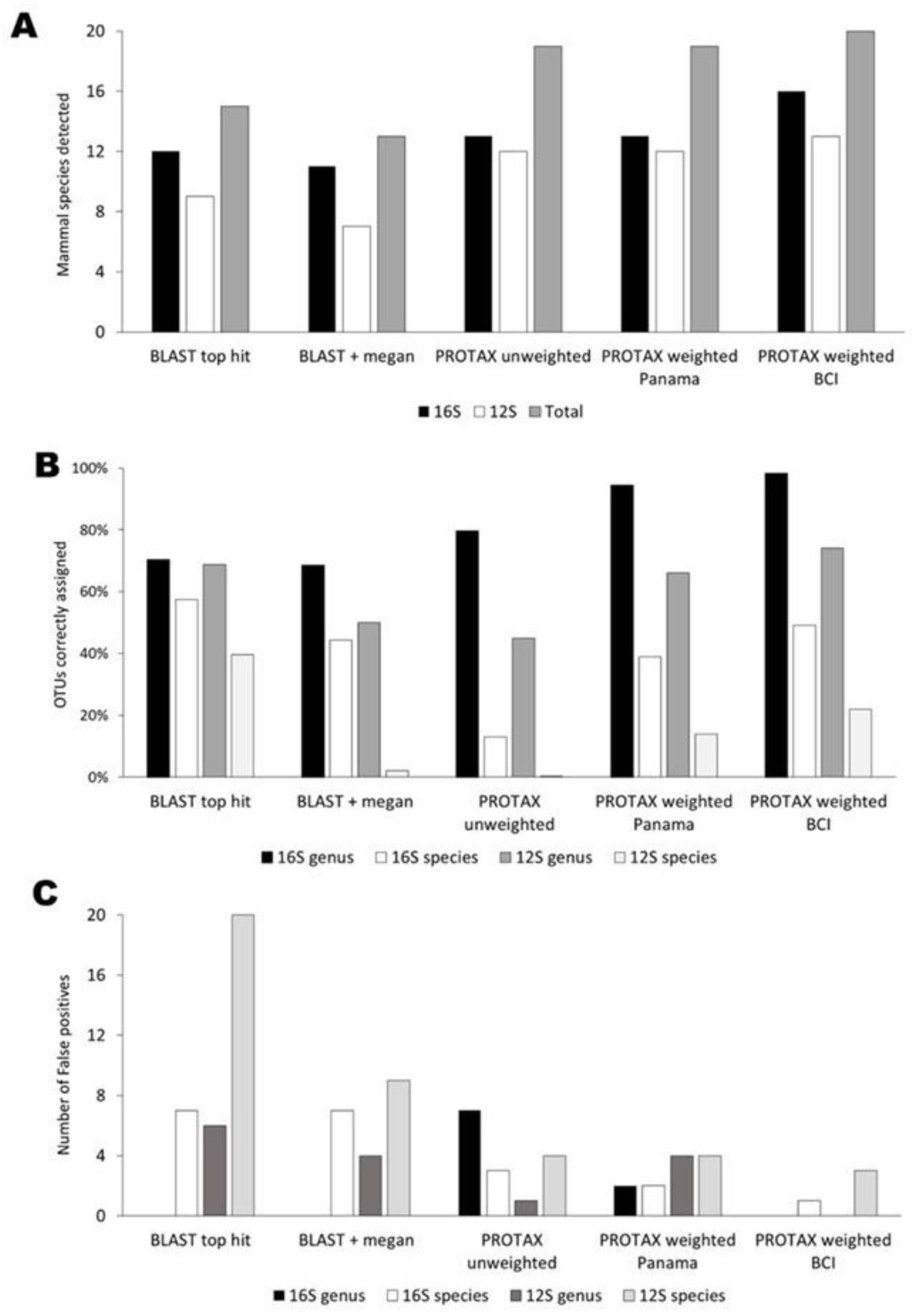
Comparison of taxonomic placement methods for assigning mammal OTUs to species from metabarcoding of carrion flies collected on Barro Colorado Island (BCI). **A**) Number of mammal species detected from BCI using each taxonomic method. **B**) Percentages of OTUs assigned to genera or species known to occur on BCI. **C**) Numbers of OTUs assigned to genera or species known not to exist on BCI (clear false positives).

**Table 1.** Twenty mammal species detected from metabarcoding of carrion flies on Barro Colorado Island Panama in 2015, along with PROTAX (BCI-weighted) estimated probabilities of correct assignment at genus and species rank, and the percentage of fly pool samples that each species was detected from.

| Species name | Common name | Genus prob. | Species prob. | % of samples |
| --- | --- | --- | --- | --- |
| *Alouatta palliata* | mantled howler monkey | 0.996 | 0.996 | 98% |
| *Ateles geoffroyi* | Geoffroy's spider monkey | 0.983 | 0.983 | 85% |
| *Tamandua mexicana* | northern tamandua | 1.000 | 0.997 | 29% |
| *Cebus capucinus* | white-faced capuchin | 0.997 | 0.979 | 26% |
| *Caluromys (derbianus)\** | Derby's woolly opossum | 0.993 | NA | 22% |
| *Saguinus (geoffroyi)\** | Geoffroy's tamarin | 0.400 | NA | 21% |
| *Sciurus granatensis* | red tailed squirrel | 0.994 | 0.994 | 21% |
| *Proechimys semispinosus* | Tome's spiny-rat | 0.457 | NA | 20% |
| *Marmosa robinsoni* | Robinson's mouse opossum | 0.982 | 0.98 | 19% |
| *Procyon (cancrivorus)\** | Crab-eating raccoon | 0.934 | NA | 16% |
| *Potos flavus* | Kinkajou | 0.979 | 0.978 | 11% |
| *Didelphis marsupialis* | common opossum | 0.984 | 0.957 | 11% |
| *Pecari tajacu* | Collared peccary | 0.979 | 0.978 | 6% |
| *Leopardus (pardalis)\** | Ocelot | 0.817 | NA | 3% |
| *Pteronotus (gymnonotus or parnellii)\** | big naked-backed bat or Parnell's mustached bat | 0.230 | NA | 3% |
| *Canis\** | Domestic dog or coyote | 0.942 | NA | 1% |
| *Bradypus variegatus* | brown-throated sloth | 1.000 | 0.998 | 1% |
| *Choloepus hoffmanni* | Hoffmann's two-toed sloth | 1.000 | 0.999 | 1% |
| *Dasyprocta punctata* | Agouti | 0.958 | 0.924 | 1% |
| *Tonatia (saurophila)\** | Stripe-headed round-eared bat | 0.894 | NA | 1% |

\* PROTAX assignment only at genus level, thus species probabilities were not estimated. Species names in parentheses are the species of the identified genus that occurs on BCI

BLAST top hit and BLAST plus MEGAN both resulted in detection of fewer mammal species (15 and 13 respectively). The percentage of OTUs assigned to a species known from BCI was higher for PROTAX than for BLAST (Fig 2b). Rates of mammal OTUs assigned to species not present on BCI (clear false positives) were also generally lower for PROTAX than for BLAST. Some clear false positives were still present with all assignment methods, although there were fewer in the weighted PROTAX analyses, and no false positives at the genus level occurred with BCI-weighted PROTAX (Fig 2c).

#### Effect of fly pooling

From the 24 flies that were extracted both singly and in pools, 3 total mammal species were detected from the pooled samples, whereas 8 total mammal species were detected from the same 24 flies when extracted singly. From the single fly extractions, a range of 1-4 species were detected from each fly.

#### Non-mammal species detection

The 12S primers also detected several birds and one lizard. We were only able to assign one OTU to the species level, wattled jacana (*Jacana jacana*). Two OTUs were assigned to birds at the genus level, one to an antshrike species (genus *Thamnophilus,* likely *atrinucha* or *doliatus;* both known from BCI), and one to a trogon species (genus *Trogon,* of which 5 species are known from BCI). Several OTUs were assigned the family Anatidae; however, these were a 100% match to many species within that family, and six species from Anatidae are known from BCI. We also assigned one reptile OTU to the whiptail lizards (family Teiidae) which is likely either *Ameiva festiva* or *Ameiva leptophrys*, the two Teiidae species known from BCI.

### Methods comparison

In 2015, we detected a greater number of mammal species in 29 days of carrion-fly metabarcoding than were detected by either 29 days of transect counts or 84 calendar days of camera trapping carried out concurrently (metabarcoding = 20 species; camera trapping = 17 species, transect counts = 13 species, Fig 3). We detected more mammal species with fly metabarcoding in 2015 than in 8 total years of transect counts from 2008-2015, but fewer than 8 years of camera trapping (transect counts 2008-2015 = 17; camera trapping 2008-2015 = 26). Using all three methods combined, we detected a total of 27 mammal species in 2015, and 34 mammal species from 2008-2015. Visual inspection of the species accumulation curves from 2015 (Fig 4) suggest that additional carrion fly sampling effort would likely have resulted in a greater number of species detections, whereas the transect count and camera-trap datasets were nearing asymptotes.

**Figure 3.**
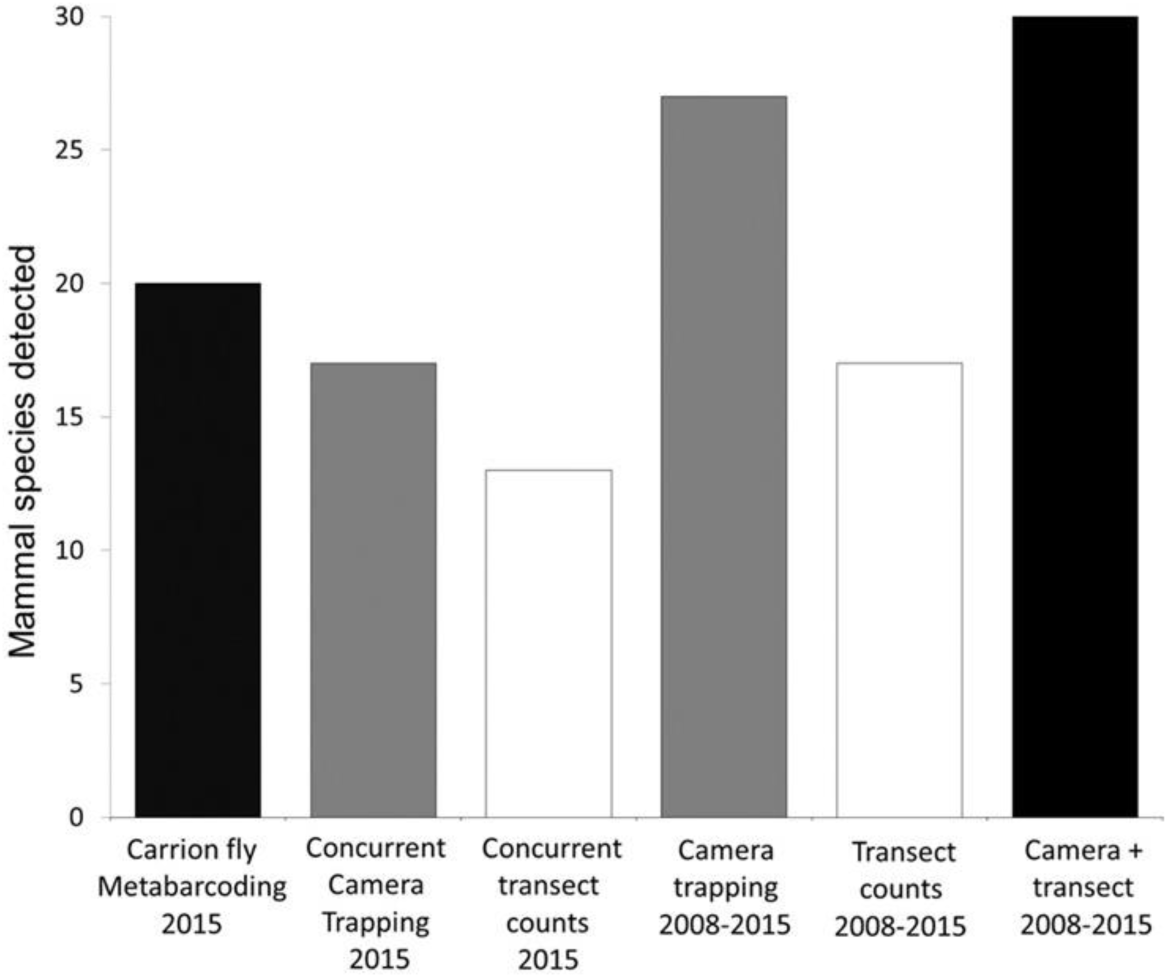
Number of mammal species detected by alternative sampling methods on Barro Colorado Island, Panama.

**Figure 4.**
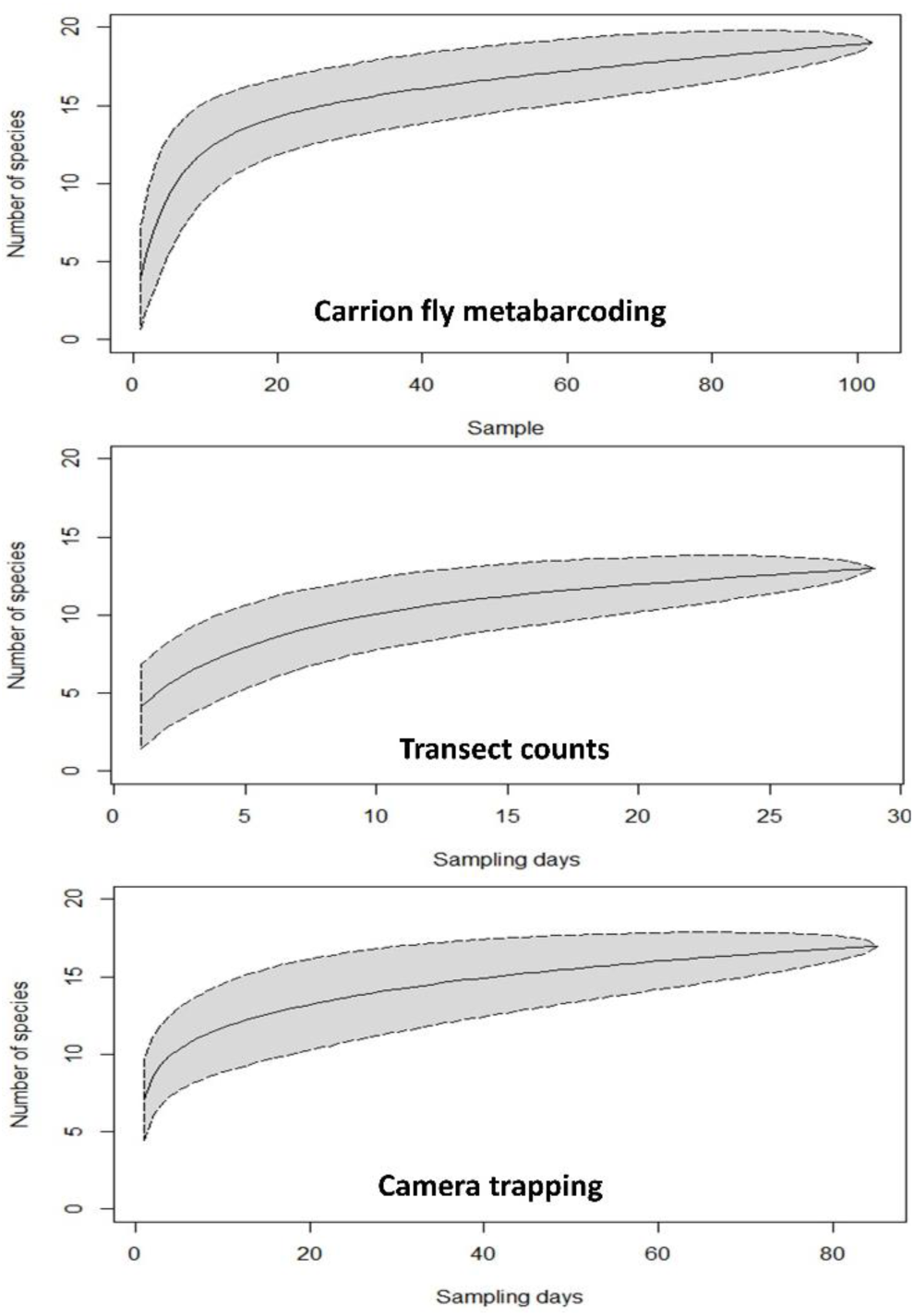
Species accumulation curves for three different survey methods used to sample mammal diversity on Barro Colorado Island, Panama in 2015.

Of the 20 species detected with fly metabarcoding (table 1), four were not detected by camera traps or transect counts from 2008-2015. These included Derby’s woolly opossum (*Caluromys derbianus*), a species from the genus *Canis* that was a 100% match to reference sequences of domestic dog (*C. domesticus)* and coyote (*C. latrans*) and two bat species. Conversely, metabarcoding did not detect three species commonly detected with transect counts or camera trapping: paca (*Agouti paca*), white-nosed coati (*Nasua narica*), and red Brocket deer (*Mazama americana*) (Table 2). Moreover, the Central-American agouti (*Dasyprocta punctata*), a large rodent that is by far the most common mammal on BCI, was detected in just one metabarcoding sample, and only by the 12S marker.

**Table 2.** Mammal species detected by carrion fly metabarcoding, camera trapping, and diurnal transect counts on Barro Colorado Island, Panama. For metabarcoding, values represent number of samples in which a species was detected. For camera trapping and transect counts, values are number of individuals detected.

| Species | common name | Metabarcoding |  | Camera trapping |  | Transect counts |  | Habit<sup>1</sup> |  |
| --- | --- | --- | --- | --- | --- | --- | --- | --- | --- |
|  |  | 16s | 12s | 2015 | 2008-2015 | 2015 | 2008-2015 |  |  |
| Order Primates |  |  |  |  |  |  |  |  |  |
| *Alouatta palliata* | Mantled howler monkey | 99 | 67 |  | 6 | 315 | 1,996 | D | A |
| *Ateles geoffroyi* | Geoffroy's spider monkey | 87 | 17 |  | 4 | 32 | 136 | D | A |
| *Cebus capucinus* | White-faced capuchin | 27 | 2 | 4 | 147 | 34 | 429 | D | A |
| *Saguinus geoffroyi* | Geoffroy's tamarin | 21 |  |  |  | 17 | 94 | D | A |
| Order Carnivora |  |  |  |  |  |  |  |  |  |
| *Potos flavus* | Kinkajou |  | 11 |  | 1 |  |  | N | A |
| *Procyon cancrivorus* | Crab-eating raccoon | 2 | 14 | 8 | 52 |  |  | N | T |
| *Leopardus pardalis* | Ocelot | 3 |  | 329 | 5,801 |  | 6 | C | T |
| *Canis sp.* | Domestic dog or coyote | 1 | 1 |  |  |  |  |  |  |
| *Nasua narica* | White-nosed coati |  |  | 1,064 | 7,358 | 18 | 178 | D | T |
| *Puma concolor* | Puma |  |  |  | 6 |  |  | N | T |
| *Panthera onca* | Jaguar |  |  |  | 9 |  |  | C | T |
| *Eira barbara* | Tayra |  |  | 1 | 54 |  | 4 | D | T |
| *Herpailurus yagouaroundi* | Jaguarondi |  |  | 3 | 5 |  |  | C | T |
| Order Rodentia |  |  |  |  |  |  |  |  |  |
| *Dasyprocta punctata* | Agouti |  | 1 | 2,975 | 72,719 | 302 | 1,821 | D | T |
| *Sciurus granatensis* | Red tailed squirrel | 10 | 15 | 20 | 1,769 | 68 | 735 | D | ST |
| *Proechimys semispinosus* | Tome's spiny-rat | 5 | 18 | 114 | 1,074 | 1 | 2 | N | T |
| *Agouti paca* | Paca |  |  | 108 | 2,412 |  |  | N | T |
| *Heteromys desmarestianus* | Desmarest's spiny pocket mouse |  |  |  | **44** |  |  | N | T |
| *Coendou rothschildi* | Rothschild's porcupine |  |  |  | **3** |  |  | N | A |
| *Diplomys labilis* | Rufous Tree Rat |  |  |  |  |  | **4** | N | A |
| *Hydrochoerus hydrochaeris* | Capybara |  |  |  | **18** |  |  | N | T |
| **Order Didelphimorphia** |  |  |  |  |  |  |  |  |  |
| *Didelphis marsupialis* | Common opossum | **1** | **11** | **57** | **1,065** |  |  | N | ST |
| *Marmosa robinsoni* | Robinson's mouse opossum |  | **19** | **1** | **7** |  |  | N | ST |
| *Caluromys derbianus* | Derby's woolly opossum | **2** | **21** |  |  |  |  | N | ST |
| **Order Pilosa** |  |  |  |  |  |  |  |  |  |
| *Choloepus hoffmanni* | Hoffmann's two-toed sloth | **1** |  |  |  | **1** | **2** | C | A |
| *Bradypus variegatus* | Brown-throated sloth | **1** |  |  |  | **1** | **2** | C | A |
| *Tamandua mexicana* | Northern tamandua |  | **30** | **20** | **359** | **2** | **27** | D | ST |
| **Order Artiodactyla** |  |  |  |  |  |  |  |  |  |
| *Pecari tajacu* | Collared peccary | **6** |  | **1040** | **12,731** | **25** | **383** | C | T |
| *Mazama temama* | Central American red brocket |  |  | **304** | **4,794** | **9** | **48** | D | T |
| **Order Cingulata** |  |  |  |  |  |  |  |  |  |
| *Dasypus novemcinctus* | Nine-banded armadillo |  |  | **5** | **169** |  |  | N | T |
| *Cabassous centralis* | Northern naked-tailed armadillo |  |  |  | **2** |  |  | N | T |
| **Order Chiroptera** |  |  |  |  |  |  |  |  |  |
| *Tonatia saurophila* | Stripe-headed round-eared bat | **1** |  |  |  |  |  | N | V |
| *Pteronotus gymnonotus* or<br>*parnellii* | Big naked-backed bat or Parnell's<br>mustached bat | **3** |  |  |  |  |  | N | V |
| **Order Perissodactyla** |  |  |  |  |  |  |  |  |  |
| *Tapirus bairdii* | Baird's tapir |  |  | **2** | **54** |  | **1** | N | T |

1 Habit: D = diurnal, C = cathemeral, N = nocturnal, A = arboreal, ST = semi-terrestrial, T = terrestrial, V = volant

## Discussion

In total, we detected a greater number of mammal species with carrion fly metabarcoding than with transect counts or camera trapping carried out during the same general time-frame with similar levels of effort. Thus, carrion fly metabarcoding appears to be an effective, albeit imperfect, method for surveys of mammalian diversity. Of the 20 species detected with fly metabarcoding (table 1), four were not detected by camera-traps or transect counts. Two were bat species that are unlikely to be detected by camera traps due to their size, and are unlikely to be detected by diurnal transect counts. The third, Derby’s woolly opossum (*Caluromys derbianus*), is arboreal and nocturnal, but it has been commonly observed on BCI at night feeding on nectar of canopy flowers. The fourth and most unexpected detection was a species from the genus *Canis*, which was a 100% match to reference sequences of both domestic dog (*C. domesticus*) and coyote (*C. latrans*). Domestic dogs are not permitted on BCI, but they are present on the mainland < 1 kilometer away at the nearest point. Coyotes have never been detected on BCI, but have been expanding in Panama (Bermudez et al. 2013) and have recently been photographed by camera traps in the adjacent Soberania National Park (www.teamnetwork.org). It is possible that a fly fed on a canid on the mainland, and then flew to BCI. Little is known about flight and foraging distances for carrion flies, so this may warrant more attention as it could affect the spatial scale of sampling.

Although fly metabarcoding detected more mammal species than traditional techniques, a clear shortcoming was failure to detect three abundant species that were commonly detected by transect counts and camera trapping. This included the common rodent (*Agouti paca*), the most common carnivore (*Nasua narica)*, and a common ungulate (*Mazama americana*). Metabarcoding did, however, detect two other rodent species, red tailed squirrel (*Sciurus granatensis*), and Tome’s spiny-rat (*Proechimys semispinosus*), and two confamilial carnivores, crab-eating raccoon (*Procyon cancrivorus*), and kinkajou (*Potos flavus*) (Table 1).

There are three potential explanations for why we were unsuccessful in detecting these three common species with metabarcoding. One possible explanation is that reference sequences for those species were missing from our reference database. Only 59% and 62% of mammal species and 82% and 85% of genera known from BCI were represented in our 16S and 12S reference databases, respectively. However, all three species were present in our reference databases to varying extents: paca was represented in our database by 12S, coati was represented by both markers, but only at the genus level (South American Coati; *Nasua nasua*), and red brocket deer was represented by both markers at the species level. Additionally, PROTAX was able to place 98% of 16S OTUs, and 74% of 12S OTUs to at least the genus level (Fig 2). Thus, missing reference sequences do not seem to be a strong explanation for our failure to detect these species. However, in less-studied areas with less complete reference-sequence representation, poor reference databases are more likely to hamper species assignment, and thus investment in building reference databases should continue.

A second potential explanation for why some common species were not detected with metabarcoding is that there were mismatches in our 16S and 12S primers with binding sites for these species. This could result in failed PCR amplification even if DNA from these species was present in fly samples. For paca, all nine reference sequences on GenBank have one mismatch in the forward primer, 6 bp from the 3’ end. One mismatch is unlikely to completely prevent amplification, but could reduce PCR efficiency, especially in mixed samples with other targets. For red brocket deer, no primer mismatches were observed for 12S. For 16S, the forward primer has one mismatch with all Genbank reference sequences, but at the far 5’ end. We could not evaluate primer mismatches for white-nosed coati at the species level, but both 16S and 12S primer sets had one 5’ mismatch to the congener *Nasua nasua.* Presently available GenBank reference sequences also may not account for local sequence variants with primer mismatches.

Lastly, it is likely that fly feeding preferences may contribute to sampling bias. Although carrion flies feed on carcasses of dead animals, we suspect that the main source of eDNA is feces. Primates were the most commonly detected species, with mantled howler monkey (*Alouatta palliata*) and Geoffroy’s spider monkey (*Ateles geoffroyi*) identified in 98% and 85% of samples respectively. Monkeys, and particularly howler monkeys which feed on leaves, produce an abundant quantity of soft scat that can be easily consumed by flies, while rodents such as agoutis and pacas, and ungulates such as red-brocket deer, have smaller and harder scats that may not be as attractive to flies. It is notable that some rodent species were detected nonetheless, but this could have been from carrion, and not from feces. Further research into the source of carrion fly derived eDNA, and how this affects detection is needed. Fly metabarcoding does seem to work especially well for primates, as all four primate species present on BCI were detected, and Calvignac-Spencer et al. (2013b) also detected many primates with carrion fly metabarcoding in Cote d’Ivoire and Madagascar.

For species assignment, PROTAX, especially weighted PROTAX, outperformed other assignment methods. With weighted PROTAX, we were able to assign nearly all OTUs to genus, and most to species, especially with 16S. For those OTUs that we could only place at the genus level, all but two we were able to assign to species based on prior information of the vertebrate community on BCI as only one member of each genus is present on the island. Weighted PROTAX resulted in fewer clearly false positives (species not known from BCI) than either BLAST or unweighted PROTAX (Fig 2c). However, even weighted PROTAX produced a few false positive assignments at the species level. In cases where it is important to be conservative with species assignment, and eliminate or minimize false assignments, selecting a high probability threshold (e.g. 0.9 or 0.95) may be desirable. All but one of our false positive assignments from weighted PROTAX at the species level, and all at the genus level, had PROTAX probabilities of < 0.9 (supplementary file S2). Assigning such a cutoff value, however, is a trade-off between minimizing false positives and false negatives. We detected several species known from BCI at assignment probabilities of < 0.9 (table 1), and so with a higher probability threshold we would have considered those species as undetected even though they are present. The results from our weighted datasets suggest that one effective way to simultaneously reduce false positive (overly confident) and false negative (overly conservative) assignments in PROTAX is to use expert knowledge to assign high prior probabilities to species known to exist in the region (Fig. 2).

As evidenced by the 24 flies that we analyzed individually and in pools, pooling reduced the number of species detected. When flies were extracted individually, we detected 8 species, including ocelot (*Leopardus pardalis*), Hoffmann’s two-toed sloth (*Choloepus hoffmanni*), collared peccary (*Pecari tajacu*), red-tailed squirrel, and all 4 primates. In the 2 pooled samples from these same 24 flies, we only detected collared peccary and the two most commonly detected primate species. Hoffmann’s two-toed sloth was only detected in one of the single-fly extractions, and was not detected in any of the pooled samples. Thus if we had not extracted some flies singly, this species would have been missed entirely.

Pooling is more likely to affect detection of rare species, as their DNA may be outcompeted during PCR by more abundant DNA of common species. It is also likely that species with more primer mismatches will be outcompeted by species with greater PCR efficiency if DNA of two such species exist in the same pool. Pooling of flies however, allows for far fewer DNA extractions and individual PCRs, which substantially reduces labor and cost (up to 16 fold in this example), which may allow many more flies to be processed, ultimately increasing the number of species detected. If a pooling strategy is employed, and there are particular target species of concern for which detection is essential, it may be desirable to include species-specific primers in addition to general primers (Schubert et al. 2014). It is also possible that the high number of PCR cycles we employed led to ‘PCR runaway’, in which common amplicons became exponentially abundant in pooled samples at the expense of less-common amplicons. Thus, we advise future studies to optimize the number of PCR cycles with quantitative PCR prior to metabarcoding (Murray et al. 2015).

The use of two different markers (12S and 16S) resulted in the detection of more species than either marker alone. Thus, we recommend using multiple markers to improve species detection. The inclusion of the 12S marker allowed us to detect more mammal species, and also allowed detection of birds and reptiles. This marker however appears to have relatively poor information content for discriminating birds and reptiles at the species level. Thus if the goal is to detect non-mammalian vertebrates it may be preferable to use additional group specific markers. Multiple markers optimized for different groups or families could be run simultaneously, which should increase detection in each group. For mammals, 12S species assignment were generally lower confidence than with 16S (supplementary file S2), and a greater proportion of 12S OTUs could not be assigned at the genus or species level (Fig 2b).

Choice of bait and trap configuration may have an impact on carrion fly metabarcoding results. We chose to use pork for bait because we wanted to target flies that feed on mammals, and because a *Sus* blocking primer has been previously designed for both markers we used (Calvignac-Spencer et al. 2013b). We made an effort to keep bait separate from flies during sampling, but even with the blocking primers, *Sus* DNA was still commonly amplified. *Sus* OTUs accounted for 68% of total 16S reads and 43% of total 12S reads in our dataset. In a small exploratory study (unpublished data), we found that increasing *Sus* blocking primer concentrations to 20x as opposed to 5x used by Calvignac-Spencer et al. (2013b) led to increased detection of non-*Sus* DNA. If pork is used as bait, we recommend taking as much care as possible to reduce contact of flies with the bait to reduce contamination, and ensure enough sequencing depth so that non-*Sus* amplicons are still sequenced even if they are in the minority. Use of other baits such as chicken or fish, or commercial fly baits may mitigate these concerns somewhat, but the 12S marker will still amplify chicken or fish DNA, and the 16S marker will also amplify fish DNA (Cannon et al. 2016). We know little about how generalized carrion flies are in their feeding habits (Calvignac-Spencer et al. 2013a). Thus, mammalian bait is still preferable for surveys aimed at mammal communities.

For maximal species detection, metabarcoding, camera-trapping, and transect counts could be used simultaneously, as they are complementary. All three techniques combined detected more species than any one of the techniques alone. Due to the nature of fly sampling, conducting fly collection and transect counts simultaneously would add little additional time or cost. During this study, we visually observed two common species missed by metabarcoding, white-nosed coati and red brocket deer, as well as numerous others, while collecting flies (unpublished data). Also, if the goal was to survey biodiversity of both vertebrates and invertebrates, fly traps could be paired with other types of insect traps e.g. pitfall traps, and fly DNA and bulk DNA from traps could be amplified with invertebrate primers and included in sequencing runs (Ji et al. 2013; Yu et al. 2012).

## Conclusions

This field test confirms that carrion fly-derived DNA metabarcoding is a powerful tool for mammal biodiversity surveys, already on par with other commonly used methods. A relatively small effort (29 days of fly sampling conducted by one individual) detected the majority of the mammal species resident on BCI (excluding bats), and a greater number of mammal species than were detected by camera trapping or diurnal transect counts with similar sampling effort. Nonetheless, carrion fly metabarcoding still failed to detect several abundant species that were easily detected with other methods, and metabarcoding may not provide reliable estimates of species relative abundance (Schnell et al. 2015). Thus, we do not advocate that carrion fly metabarcoding replace current methods, but rather add to the tools available for biodiversity surveys. Our results suggest that methodological modifications would likely increase the detection power of carrion fly metabarcoding still further, including more complete reference sequence databases, optimization of PCR conditions, multiple custom markers, and moderately greater sampling effort. Individual fly metabarcoding would also increase detection power, but at a larger cost. Nevertheless, we conclude that carrion fly metabarcoding is an important tool, with the potential to greatly improve researchers’ ability to efficiently monitor biodiversity. As sequencing costs drop and as reference databases of complete mitochondrial genomes becomes readily available (Tang et al. 2014), this method promises to allow for the rapid characterization and monitoring of vertebrate communities.

## Acknowledgments

This work was funded by a Smithsonian Tropical Research Institute fellowship (TR). Camera trapping and transect counts was funded by the STRI Terrestrial Environmental Studies Project, the U.S. Department of Education Math-Science Partnerships Project (JG), and private funds from Gregory E. Willis. We also thank Gregory E. Willis for years of mammals census field work. DY was supported by the National Natural Science Foundation of China (31400470, 41661144002, 31670536, 31500305, GYHZ1754), the Ministry of Science and Technology of China (2012FY110800), the University of East Anglia, and the State Key Laboratory of Genetic Resources and Evolution at the Kunming Institute of Zoology (GREKF13-13, GREKF14-13, GREKF16-09).

## Supplementary files

**S1** Spreadsheet of weighted and unweighted PROTAX probabilities at the Class, Family, Genus, and Species level for all OTUs.

**S2** FASTA file of OTU sequences.

## Data Accessibility

Raw sequence data are available on the NCBI Sequence Read Archive under accession number PRJNA382243.OTU sequences are available in the supplementary information. Scripts for the PROTAX analysis will be made available on Dryad prior to publication.

## Author contributions

Torrey Rodgers designed the study, carried out all fly collection field work and lab work, coordinated the data analysis, and wrote the manuscript. Charles Xu performed the PROTAX analysis, Jacalyn Giacalone collected the camera trap and transect count data, Karen Kapheim conducted the sequence filtering and OTU generation bioinformatics work, Kristin Saltonstall helped with design and implementation of lab work, Marta Vargas helped with lab work management and conducted MiSeq sequencing, Douglass Yu supervised the PROTAX analysis, Panu Somervuo provided scripts and helped with the PROTAX analysis, W. Owen McMillan provided lab space and payed for sequencing, and Patrick A. Jansen helped with field study design. W. Owen McMillan and Patrick Jansen jointly advised this work. All authors contributed to manuscript editing.

